# An Overview of the Application of Deep Learning in Short Read Sequence Classification

**DOI:** 10.1101/2020.09.19.304782

**Authors:** Kristaps Bebris, Inese Polaka

## Abstract

Advances in sequencing technology have led to an ever increasing amount of available short read sequencing data. This has, consequently, exacerbated the need for efficient and precise classification tools that can be used in the analysis of this data. As it stands, recent years have shown that massive leaps in performance can be achieved when it comes to approaches that are based in heuristics, and alongside these improvements there has been an ever increasing interest in applying deep learning techniques to revolutionize this classification task. We attempt to gather up these approaches and to evaluate their performance in a reproducible fashion to get a better perspective on the current state of deep learning based methods when it comes to the classification of short read sequencing data.

## I. Introduction

Individual genome and metagenome sequencing has become increasingly more affordable over the past years [1][2], which has lead to an explosion in the amount of data that is available for analysis. This has, in turn, spawned a need for more affordable tools to analyze this data. In this study we will look at recently developed taxonomic classification tools and evaluate their performance when working with metagenome sequencing data. A state of the art tool widely used for tasks such as this is Kraken2 [3], which has shown that there is still a lot of room for improvement in currently used methods, with Kraken2 reducing its memory footprint to just 15% and its runtime to just 20% its predecessor Kraken [3].

A number of deep learning and machine learning based tools have been produced in recent years with a goal of alleviating different, resource intensive, aspects of current methods – memory requirements, classification time, classification precision or disk space requirements [4][5][6][7]. A major issue of evaluating the current state of these tools is in the fact that there are no published results showing their relative performance when the same reference data is used and when they’re applied to non-synthetic samples. This is an issue that is characteristic to many bioinformatics tools [8]. There is also a known issue with self reported performance metrics, which are the only results available for a lot of tools – they tend to have bias problems [9].

There have been some attempts to standardize the way tools are benchmarked [8], but thus far no standard protocols nor benchmark data sets have been adopted and most tools still report their results using bespoke approaches. This makes gauging the performance of these tools in real world scenarios quite tricky as generated samples can be less complex than real world ones [16].

Our goal is to evaluate the performance of a set of freely available published deep learning based tools by using the exact same reference data: metagenome data from a set of samples that have been sequenced by MGI DNBSEQ-T7 sequencers [10], which include 2 reference samples with a predefined composition (Zymo samples [11] – referred to as C1 and C2) and 3 human gut (fecal) metagenome samples that were produced within the ERDF project ‘Optimisation of H.pylori eradication therapy for population-based gastric cancer prevention’ (referred to as S1, S97 and S104). The non-control samples were added to the experiment to assess how the tools perform if the reference database does not contain all of the organisms present in the sample and therefore there are no exact matches to the sequences in the sample.

## II. Methods

We selected suitable deep learning based tools for genomic classification by searching the SCOPUS and Arxiv databases, excluding the tools that were designed for tasks other than shotgun sequence classification. This way we picked 4 tools:

- MetaVW – embedding based tool, leverages Vowpal Wabbit [12] as its backbone [4],
- fastDNA – fastText [18] based embedding tool [5],
- GeNet – a classification tool that uses a convolutional model [6],
- DeepMicrobes – a classification tool that leverages an attention model [7].

In addition, we elected to use the Kraken2 results as our ‘ground truth’ for samples that we did not have a theoretical taxonomic distribution for because it is a widely recognized and commonly used tool [3]. We found this necessesary due to the way the experiment was set up: we included real-life samples with an unknown composition, so an estimate of their composition was established using Kraken2 as a reliable state-of-art tool.

The expirements were performed on a workstation running a 12 core procesor (Intel i7-8700k), 64 GB of RAM, a Nvidia GTX 1080 Ti graphics card (GPU) and 500 GB of PCIe storage. We make a point of noting down the storage solution due to Vowpal Wabbit being heavily bound by storage speed [12] which will impact the performance of MetaVW [4].

We evaluated all tools using the same 5 samples (read counts and average read length, which provide insight into data complexity, are given in Table 1) and used the small database from MetaVW [13] (mostly due to practical concerns – it contained all of the info needed for all of the tools to train) to create the reference databases for all of the tools (containing 1565 sequences related to 193 species) [4].

**Table I.** Sample characteristics

| Sample | Forward read count | Forward average read length | Reverse read count | Reverse average read length |
| --- | --- | --- | --- | --- |
| **C1** | 38581675 | 150 | 38581675 | 150 |
| **C2** | 52537916 | 150 | 52537916 | 150 |
| **S1** | 28990760 | 150 | 28990760 | 150 |
| **S97** | 28987851 | 150 | 28987851 | 150 |
| **S104** | 22358511 | 150 | 22358511 | 150 |

The two control samples are ZymoBIOMICS™ Microbial Community Standard samples and contain data from 10 different species [11].

Reference databases were created and models were trained using the default setting wherever possible. The only deviation from this being DeepMicrobes where we used 11-mers instead of 12-mers due to not being able to train the 12-mer model within 11 GB of video RAM. The commands we used have been made available on GitHub [14]. These runs were timed and resource usage was monitored during training. A general outline of the database creation process is presented in Figure 1.

**Fig. 1.**
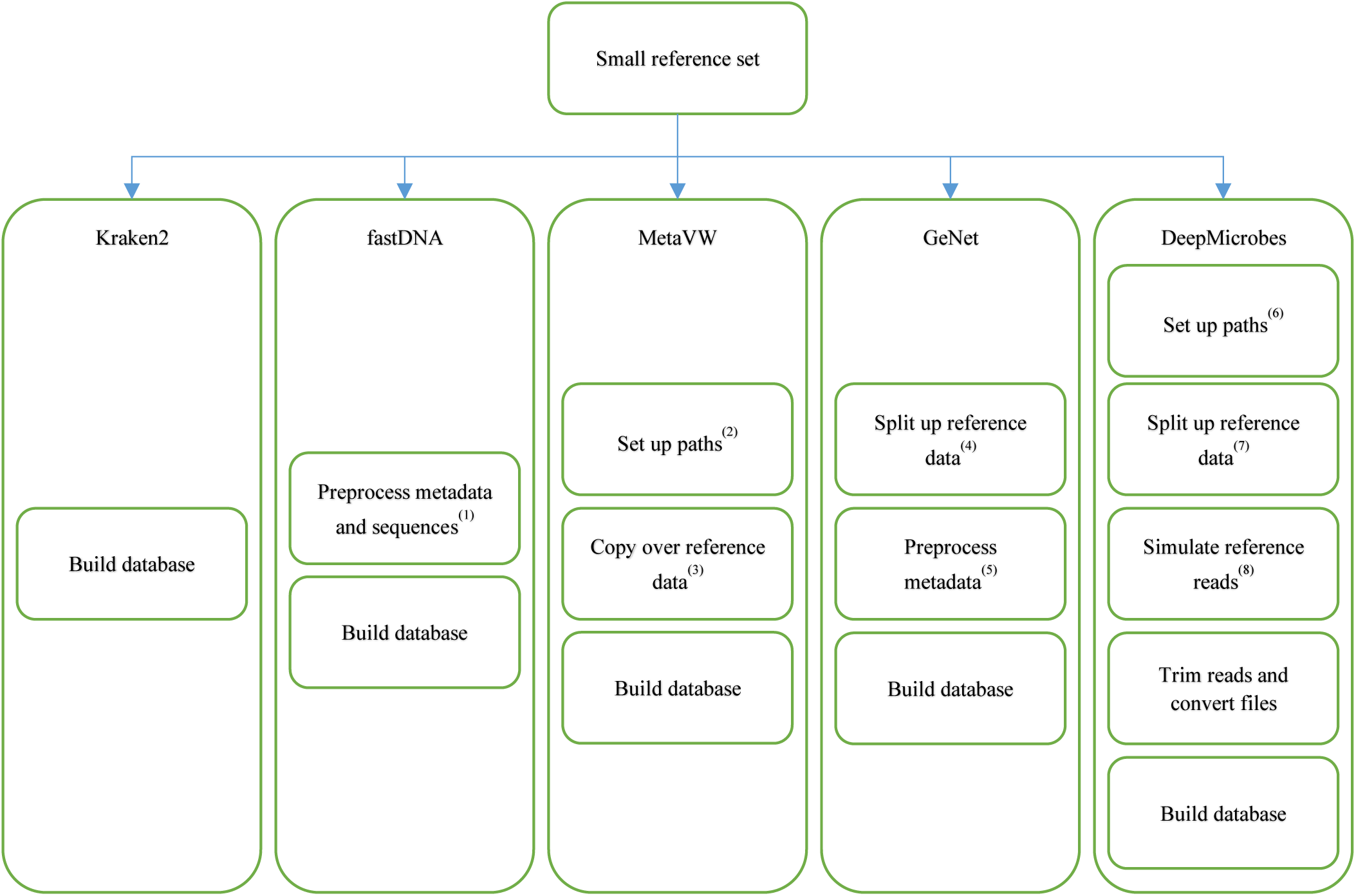
An overall outline of the database creation process. Some practical items that we would like to draw additional attention to include: fastDNA (1) requires all sequences to be uppercase and needs custom meta tags for the sequences, MetaVW (2) requires some external libraries that the tool does not ship with, GeNet needs the reference data to be split up on a taxon-by-taxon basis and (4) requires custom meta tags for the sequences, DeepMicrobes (5) ships with all of the required tools, but they’re split up between a multitude of subdirectories, (6) it also requires the reference data to be split up into separate files similarily to GeNet and (7) the training set has to be processed with a read simulator [18]. The scripts that this figure represents are available on GitHub [14].

After training the neural network models, they were used to classify the previously described samples. The results were evaluated using both the necessary computing resources, which were quantified as runtime and resource usage data (memory and graphics card usage), and the classification precision, based on the ‘ground truth’: the theorethical composition of the reference control samples or composition identified by Kraken2 for the gut microbiome samples. We calculated the overall precision as the proportion of reads that the current tool has classified the same as the ‘ground truth’:

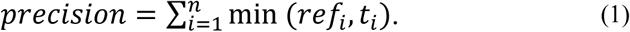

We also evaluated the overall coverage, which was achieved by each of the tools. The coverage is a relevant metric due to its ability to clearly show if a tools precision could be an artefact of it either under or over reporting when compared to other tools and was calculated as:

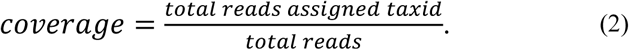

Classification results (precision and coverage) were evaluated both in raw form, where the resulting taxonomic rank of the organism in the taxonomy tree was defined by the tool, and normalized in 2 different ways to determine the suitability of produced results for practical applications: selecting the results classified at the phylum rank or at the genus rank. This was done by traversing taxonomic tree data that was aquired from NCBI [16].

## III. Results and Discussion

Overall training process showed a high dispersion in terms of time to train/build. Kraken2 and fastDNA showed the best results (Table 2). In all probability fastDNA outperformed the rest of the tools that leverage neural networks by such a large margin due to it being based on the highly efficient fastText [18]. Interestingly, there is a significant difference between GeNet and DeepMicbrobes and between MetaVW and fastDNA, even though the two pairs use similar methods when approaching the problem of learning the contents of the reference set. The disparity between MetaVW and fastDNA could be explained by the multithreaded capabilities of the tools that they are based on [12][17]. While the disparity between GeNet and DeepMicrobes comes down to termination conditions – with DeepMicrobes terminating after 1 epoch by default [19] while the implementation of GeNet implies that it should terminate either at 400 epochs or when reaching 99.5% precision, but it did not – we stopped the training process by hand once it had reached 99.5% precision (at epoch 418) [19].

**Table 2.** Training performance

| Tool | Memory Usage | Time taken (including preprocessing) | Is GPU accelerated |
| --- | --- | --- | --- |
| **Kraken2** | 2939 MB | 3m (16m) | No |
| **MetaVW** | 15.7 GB | 15h47m | No |
| **GeNet** | 2964 MB | 10d23h50m | Yes |
| **DeepMicrobes** | 5958 MB | 6h15m (6h39m) | Yes |
| **fastDNA** | 3205 MB | 16m | No |

It should be noted that a signicifant portion of the training time for DeepMicrobes was spent on preprocessing the samples for further use. This along with the time that was necesary for Kraken2 to prepare for database being built has been separated out to give a better idea of the time these two tools spends on the build/train process itself (Table 2).

Choosing Kraken2 as the baseline when data on sample composition were not available carries a caveat: accuracies for the non-control samples are not necessarily indicative of the best performance – they are indicative of the performance most similar to Kraken2.

Some concessions had to be made when performing the classification tasks. We excluded GeNet from the classification tasks since it had no readily made classification scripts and producing such scripts was out of the scope of this article. MetaVW and fastDNA required single input files (we found functions that implied that paired end data could be fed into fastDNA, but could not find a way of accessing it without modifying the tool [20]), which meant that samples were merged using bbmerge. Additionally, DeepMicrobes could process controls as is, but required additional utility scripts to be produced to facilitate working with non-control samples due to the large amount of storage space that preprocessing each sample required (a 9 GB gzipped sample expanded to 127 GB in temporary files that resulted in a 86 GB tfrec file). Therefore we batched the samples into random 10% subsamples whose results were then joined; as a result the reported runtimes are based solely on the controls.

Just like when training the model preprocessing samples for DeepMicrobes took a significant amount of time. It is indicated separately in the table in brackets (Table 3).

**Table 3.** Test performance

| Tool | Memory Usage | Time taken/sample | Is GPU accelerated |
| --- | --- | --- | --- |
| **Kraken2** | 2104 MB | 27s | No |
| **MetaVW** | 8204 MB | 23m | No |
| **GeNet** | - | - | - |
| **DeepMicrobes** | 2882 MB | 5h36m (8h14m) | Yes |
| **fastDNA** | 3207 MB | 24s | No |

When it comes to the amount of reads that each of the tools reported as classified, DeepMicrobes and MetaVW differentiated themselves from the rest by not reporting any results as unclassified, but always reporting some taxon ID. FastDNA fell on the other end of the spectrum classifying around 8% of the reads. We found Kraken2 to be the only tool where the number of reads that the tool was able to classify was related to the complexity of the sample at hand: the results for control samples composed of few different species were much higher than for the diverse human samples (Table 4).

**Table 4.** Coverage

| Tool | C1 | C2 | S1 | S97 | S104 |
| --- | --- | --- | --- | --- | --- |
| **Kraken2** | 84.94% | 82.38% | 7.95% | 6.93% | 5.24% |
| **MetaVW** | 100% | 100% | 100% | 100% | 100% |
| **GeNet** | - | - | - | - | - |
| **DeepMicrobes** | 100% | 100% | 100% | 100% | 100% |
| **fastDNA** | 8.69% | 8.60% | 8.32% | 8.43% | 8.47% |

We found that MetaVW outperforms every other tool (even Kraken2) when examining the precision metric of the baseline results (Table 5). It should be noted that these results are skewed as both Kraken2 and DeepMicrobes traverse the taxonomic tree when they are not confident enough to classify something at a certain taxonomic rank of the said tree [3][7]. But it is still noteworthy that MetaVW can be reasonably used to gather insight from simple samples without the need for normalization scripts.

**Table 5.** Precision

| Tool | C1 | C2 | S1 | S97 | S104 |
| --- | --- | --- | --- | --- | --- |
| **Kraken2** | 33.48% | 27.25% | * | * | * |
| **MetaVW** | 66.68% | 52.27% | 1.69% | 1.33% | 1.16% |
| **GeNet** | - | - | - | - | - |
| **DeepMicrobes** | 0% | 0% | 0.07% | 0.05% | 0.04% |
| **fastDNA** | 1.01% | 1.02% | 92.13% | 92.08% | 91.86% |

\*Used as reference (100%).

**Table 6.** Coverage Genus

| Tool | C1 | C2 | S1 | S97 | S104 |
| --- | --- | --- | --- | --- | --- |
| **Kraken2** | 84.21% | 81.49% | 4.02% | 3.36% | 2.54% |
| **MetaVW** | 99.84% | 99.83% | 88.39% | 90.52% | 92.45% |
| **GeNet** | - | - | - | - | - |
| **DeepMicrobes** | 70.33% | 77.64% | 61.66% | 60.83% | 59.03% |
| **fastDNA** | 8.56% | 8.65% | 8.28% | 8.39% | 8.43% |

When data is normalized to genus (that is, all of the items above genus are brough down to genus rank – all others are marked as unclassified), we see some interesting patterns emerge: the reported percentages of most tools drop, with Kraken2 dropping around half of the classified reads for the non-control samples and DeepMicrobes dropping around 25% of the reads.

This increases the overall classification precision of Kraken2 up to around the performance of MetaVW with the other two tools underperforming by more than 60 percentage points when it comes to the controls. When looking at the non-control samples, interestingly, MetaVW drops below both DeepMicrobes and fasDNA in terms of precision. This can mainly be explained by the large amount of unclassified reads present in the Kraken2 results. It is interesting to note that fastDNA performs very closely to Kraken2 when looking at the non-control samples (Table 7).

**Table 7.** Precision Genus

| Tool | C1 | C2 | S1 | S97 | S104 |
| --- | --- | --- | --- | --- | --- |
| **Kraken2** | 66.66% | 47.98% | * | * | * |
| **MetaVW** | 67.79% | 54.41% | 15.52% | 12.84% | 10.09% |
| **GeNet** | - | - | - | - | - |
| **DeepMicrobes** | 6.18% | 12.00% | 38.56% | 39.38% | 41.14% |
| **fastDNA** | 2.38% | 2.41% | 93.03% | 93.27% | 92.68% |

\*Used as reference (100%).

Normalizing to phylum helps most of the tools retain a larger number of reads as expected since phylum is a rank that includes genus [21]. The reduction in coverage is less than one percentage point for all tools except DeepMicrobes (Table 8).

**Table 8.** Coverage Phylum

| Tool | C1 | C2 | S1 | S97 | S104 |
| --- | --- | --- | --- | --- | --- |
| **Kraken2** | 84.92% | 82.34% | 7.85% | 6.84% | 5.16% |
| **MetaVW** | 100% | 100% | 100% | 100% | 100% |
| **GeNet** | - | - | - | - | - |
| **DeepMicrobes** | 74.23% | 80.07% | 67.81% | 67.45% | 67.96% |
| **fastDNA** | 8.60% | 8.69% | 8.32% | 8.43% | 8.47% |

The pattern that the tools fall into does not differ from the results that we saw when normalizing for genus: precision improves significantly for less diverse samples and there is some improvement from baseline for other samples, although the increase is not as significant as in genus data (Table 9).

**Table 9.** Precision Phylum

| Tool | C1 | C2 | S1 | S97 | S104 |
| --- | --- | --- | --- | --- | --- |
| **Kraken2** | 84.91% | 72.12% | * | * | * |
| **MetaVW** | 92.59% | 80.20% | 7.85% | 6.84% | 5.16% |
| **GeNet** | - | - | - | - | - |
| **DeepMicrobes** | 36.00% | 36.00% | 33.88% | 34.22% | 33.49% |
| **fastDNA** | 6.37% | 6.43% | 96.26% | 95.34% | 95.25% |

\*Used as reference (100%).

The coverage of DeepMicrobes diminished significantly during normalization and this pattern was not observed in other tools. Therefore we investigated the taxonomic division of the results for all of the tools. fastDNA provided species results, MetaVW mostly provided species results as well as sometimes responded with taxon IDs that corresponded to genotype (less than 1% of the reads). Kraken2 traversed most of the tree with all returned taxids being valid: Kraken2 classified around 37% of the control sample reads and around 1.4% of the non-control sample reads as species. We found that unlike Kraken2 DeepMicrobes can generate arbitrary taxon IDs that cannot be found in the NCBI taxonomy data: overall DeepMicrobes returned taxon IDs that correspond to species for around 50% of the reads and classified around 20% of the control reads, while 30% of the non-control reads had been assigned non-existant taxon IDs [14]. The prevalence of these non-existant taxon IDs strongly implies that the training set we used is too small for DeepMicrobes to function properly.

## IV. Conclusions

We found that tools that leverage deep learning still have some catching up to do when comparing their performance to Kraken2 in terms of both memory usage and runtime. With Kraken2 using less than a fifth of the memory of the most memory intensive tool and undercutting some of the tools more than 500 times when looking at runtimes at the scale that we tested at. A notable exception when looking at the runtimes is fastDNA – its performance was comparable to Kraken2. It was interesting to see that embedding based tools perform so similarily to Kraken2 when evaluating their coverage and classification precision, even if this similarity in performance is limited to specific types of samples depending on the thresholding strategy that the tool employs. While the performance of DeepMicrobes did not meet the prior expectations (based on results reported in the literature) and we did not manage to fully benchmark GeNet, we still see a lot of potential in using such tools in environments where system memory is limited and researchers are willing to wait for a longer amount of time to obtain the results. It is important to note that this is a statement that heavily hinges on the comparative performance of these tools when they are trained with a suitably large reference set.

When looking at the results, we can see that the most useful and operable tool is still Kraken2, but there is a disadvantage – using Kraken2 with a larger database still requires a significant amount of RAM (around 200 GB [22]). This is where the deep learning based tools should readily outscale Kraken2.

We believe that a different thresholding mechanism for fastDNA could possibly make it competitive with Kraken2 even when using a small reference database such as the one that was used in this study. And that exploring the behaviour of these tools with significantly larger reference databases could yield more insights into how well these tools scale with the amount of available data.

Overall we would like to conclude that a lot of promising progress has been made when it comes to applying deep learning to bioinformatics and we are eager to see what the future will bring. We are hopeful that this paper draws more interest to these tools and applying deep learning to bioinformatics. Additionally we hope for someone with more complex control samples to evaluate the performance of these tools.

An interesting byproduct of performing this study is the realization that these tools are harder to use than we had initially expected. We attempt to make replicating this study easier by outlining the setup process in the repository that contains the scripts that were used to obtain these results. This has been made publicly available on GitHub [14].

## V. Acknowledgements

We would like to thank the Institute of Clinical and Preventitive Medicine of the University of Latvia and Latvia MGI Tech for gathering and sequencing the samples that made this article possible. The samples were sequenced within the ERDF project No. 1.1.1.1/18/A/184 ‘Optimisation of H.pylori eradication therapy for population-based gastric cancer prevention’.

**Kristaps Bebris** currently holds a Masters degree with a specialization in bioinformatics from Uppsala University from which he graduated in 2017. He is currently a research assistant at the University of Latvia, data scientinst at Sannsyn Latvia and a PhD student at Riga Technical University, His main interests are natural language processing, bioinformatics and genetic algorithms.

**Inese Polaka** received her Doctoral degree (Dr.sc.ing.) in information technology from Riga Technical University in 2014. She has since worked at Riga Technical University and holds the position of Assistant Professor at the moment. Her fields of interest are machine learning, data mining, especially in medical applications, evolutionary algorithms, transparent and interpretable supervised and unsupervised learning methods, as well as bioinformatics (main focus on prokaryotic metataxonomics and metagenomics).

